# Genetic sensitivity analysis: adjusting for genetic confounding in epidemiological associations

**DOI:** 10.1101/592352

**Authors:** Jean-Baptiste Pingault, Frühling Rijsdijk, Tabea Schoeler, Shing Wan Choi, Saskia Selzam, Eva Krapohl, Paul F. O’Reilly, Frank Dudbridge

## Abstract

Associations between exposures and outcomes reported in epidemiological studies are typically unadjusted for genetic confounding. We propose a two-stage approach for estimating the degree to which such observed associations can be explained by genetic confounding. First, we assess attenuation of exposure effects in regressions controlling for increasingly powerful polygenic scores. Second, we use structural equation models to estimate genetic confounding using heritability estimates derived from both SNP-based and twin-based studies. We examine associations between maternal education and three developmental outcomes – child educational achievement, Body Mass Index, and Attention Deficit Hyperactivity Disorder. Polygenic scores explain between 14.3% and 23.0% of the original associations, while analyses under SNP- and twin-based heritability scenarios indicate that observed associations could be almost entirely explained by genetic confounding. Thus, caution is needed when interpreting associations from non-genetically informed epidemiology studies. Our approach, akin to a genetically informed sensitivity analysis can be applied widely.

**Author summary:** An objective shared across the life, behavioural, and social sciences is to identify factors that increase risk for a particular disease or trait. However, identifying true risk factors is challenging. Often, a risk factor is statistically associated with a disease even if it is not really relevant, meaning that even successfully improving the risk factor will not impact the disease. One reason for the existence of such misleading associations stems from genetic confounding. This is when genetic factors influence both the risk factor and the disease, which generates a statistical association even in the absence of a true effect of the risk factor. Here, we propose a method to estimate genetic confounding and quantify its effect on observed associations. We show that a large part of the associations between maternal education and three child outcomes - educational achievement, body mass index and Attention-Deficit Hyperactivity Disorder-is explained by genetic confounding. Our findings can be applied to better understand the role of genetics in explaining associations of key risk factors with diseases and traits.

## Introduction

Associations between exposures and outcomes are commonly reported in epidemiological research, but often without estimating or accounting for the contribution from genetics. However, most exposures and outcomes are substantially heritable, and genetics can confound these associations. Here, we propose a new genetic sensitivity analysis, which we call *Gsens*, to assess to what extent genetic confounding can account for observed associations.

### Genetic confounding and sensitivity analysis

Identifying exposures that can be targeted in effective interventions is a fundamental objective shared across the life, behavioural and social sciences. To this end, identifying *causal* exposures is essential as interventions that target non-causal exposures will likely fail. To establish causation, it is necessary to account for confounding, which happens when a third variable causally influences both the exposure and the outcome, thereby generating a non-causal association between them. Genetic confounding is a special case when genetic factors play the role of the third variable. The concept of genetic confounding was introduced during the controversy regarding the effect of cigarette smoking on lung cancer. In a letter entitled 'alleged dangers of cigarette-smoking', Ronald Fisher qualified ’the mild and soothing weed’ as ‘possibly an entirely imaginary cause’ for lung cancer [1]. He argued that genetic factors could directly influence both smoking and lung cancer, generating a non-causal association between them. Although Fisher was mistaken in this particular instance, the notion of genetic confounding remains relevant, in his words ‘a common cause, in this case the individual genotype’. During this controversy, Jerome Cornfield argued against this ‘constitutional hypothesis’ [2,3]. He contended that implausibly large genetic effects (or other unobserved confounders) would be required to explain away all of the observed association. This led to the birth of the approach now called *sensitivity analysis*, which consists of estimating how strong an unknown confounder needs to be in order to explain away an observed association, providing insights into the robustness of that association (i.e. how sensitive it is to confounding and whether it is likely causal or not) [2]. Since then, sensitivity analyses have become common epidemiological tools to probe the robustness of findings under alternative scenarios. However, sensitivity analysis using genetic data has not progressed. We recently [4] proposed to use polygenic scores – individual-level scores that summarize genetic risk (or protection) for a given phenotype – to estimate the proportion of observed associations explained by genetic confounding. However, because polygenic scores capture only a small part of heritability, controlling for polygenic scores cannot entirely capture genetic confounding. We therefore propose a sensitivity analysis using polygenic scores to gauge how likely it is that genetic confounding accounts, in part or entirely, for a given exposure-outcome association. Here, we develop this proposition in two stages. First, we test to what extent associations of interest are accounted for by observed polygenic scores. Second, in the sensitivity analysis per se, we use structural equation models to examine how an increase in the predictive power of polygenic scores based on heritability estimates would affect association estimates. This can be thought of as adjusting for latent polygenic scores that capture as much of the variance in the exposure and outcome as suggested by available heritability estimates.

### Maternal education and child developmental outcomes

To illustrate our approach, we focus on maternal educational attainment (termed maternal education) as the exposure of interest. Greater maternal education is associated with child developmental outcomes in several key domains: social development (e.g. better educational outcomes), physical health (e.g. lower Body Mass Index, BMI), and mental health (e.g. lower levels of Attention-Deficit Hyperactivity Disorder (ADHD) symptoms)[5–8]. However, observed associations between maternal education and developmental outcomes are not free from confounding, in particular genetic confounding as both maternal education and developmental outcomes are heritable, and mother and child share half their genomes identical by descent [5,9–13].

Here, we illustrate the use of *Gsens* to estimate the role of genetic confounding in explaining the associations between maternal educational and three developmental outcomes in the child: educational achievement operationalized by the General Certificate of Secondary Education (GCSE), BMI, and ADHD. Importantly, *Gsens* has a wide scope of applications as it only requires genome-wide data on large samples and a focus on outcomes for which polygenic scores are available. Its applicability will further expand with the steady increase in the number and the accuracy of available polygenic scores [14].

## Results

### Method Overview

Participants were drawn from the Twins Early Development Study (TEDS), with sample sizes between 3,663 and 4,693 individuals with data for maternal education and child educational achievement, BMI, and ADHD. Polygenic scores were estimated in the child using PRSice software [15] at different p-value thresholds, explaining increasing amounts of variance in the corresponding phenotype. In the first stage, we estimated the proportion of the observed phenotypic association between the exposure and the outcome that was explained by polygenic scores at different p-value thresholds; we call these the *observed scenarios*. However, even the best-fitting polygenic scores only captured a fraction of the heritability of their corresponding phenotypes, thus underestimating the magnitude of genetic confounding. In the second stage, the sensitivity analysis therefore aimed to answer the following question: to what extent is the exposure X associated with the outcome Y after controlling for all genetic confounding? In other words, if *β_XY_* is the coefficient of regression of Y on X, to what extent would it attenuate if we were to control for ‘perfect’ polygenic scores capturing all genetic influences on X and Y rather than the small fraction accounted for by available polygenic scores? To this end, we estimated *β_XY_* under plausible scenarios combining information on current polygenic scores and heritability estimates. The estimation of *β_XY_* is based on the matrix of observed correlations between polygenic scores, exposure and outcomes. We then fit a Structural Equation Model to this matrix of correlations that aims to reflect the true extent of genetic confounding (see Methods). Approaches using one polygenic score (for the exposure or for the outcome) or two polygenic scores (for the exposure and the outcome) were used. Three functions are provided that adjust the association of interest based on the polygenic score for the exposure (GsensX), for the outcome (GsensY) or both exposure and outcome (GsensXY). We conducted simulations to assess the relative accuracy of these functions and to assess the effect of unobserved non-genetic confounding on the estimates obtained from *Gsens*. We provide a package and a tutorial at https://github.com/JBPG/Gsens.

### Observed and heritability-based scenarios

As shown in Table 1, the best-fitting polygenic scores derived from the GWAS for years of education, BMI and ADHD explained a substantial amount of the variance of, respectively, child educational achievement (threshold of p = .158), BMI (threshold: p = .20) and ADHD symptoms (threshold: p = 0.358) in TEDS. All three were highly significant (largest p value = 1.6e-20 for ADHD). Table 1 shows parameters for two main heritability-based scenarios: SNP-based and Twin-based heritability. SNP-based heritability estimates were obtained through LD score regression [16,17], based on LD Hub [18] for years of education and BMI and the most recent ADHD GWAS for ADHD [13]. Twin-based estimates were derived from TEDS and from the literature (see Table 1 note).

**Table 1.**
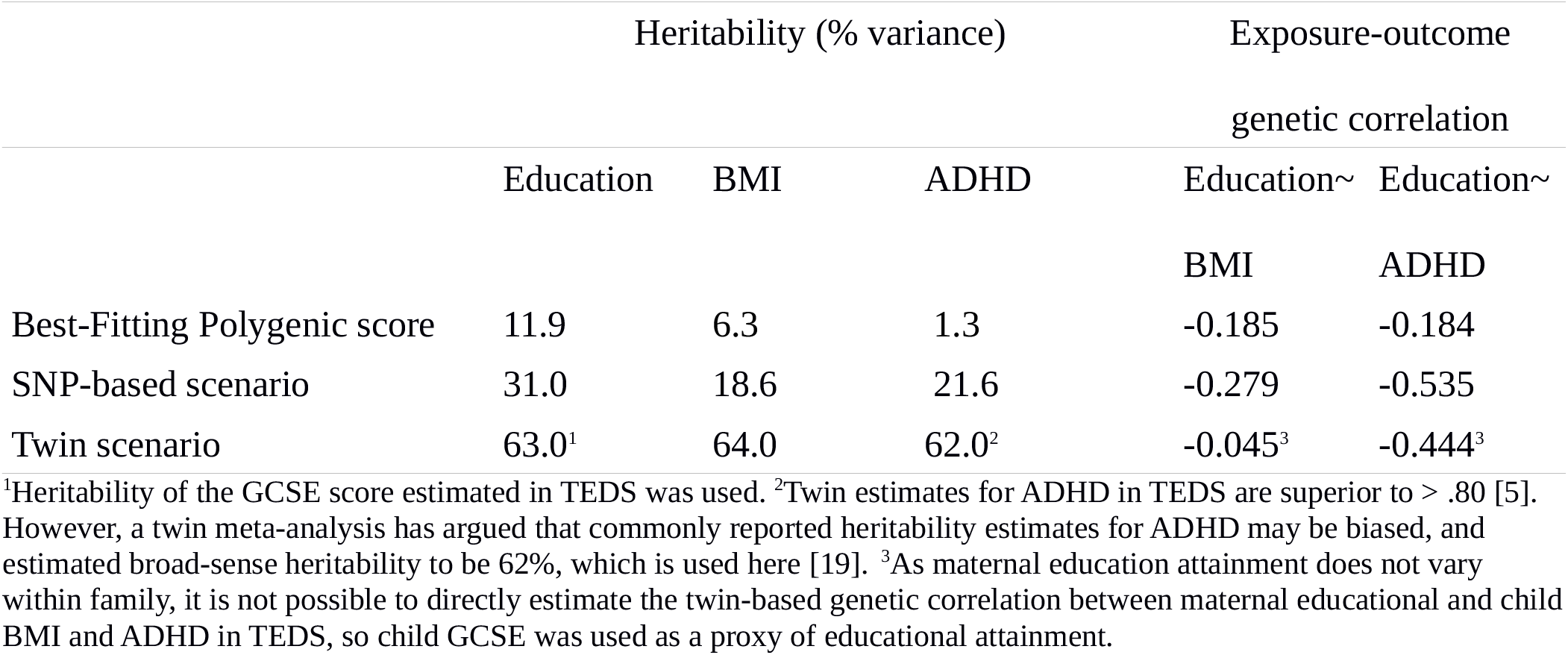
Heritability and genetic correlation under different scenarios.

### Genetic confounding and sensitivity analyses

#### Single polygenic score: child educational achievement

Supplementary material S1 eTable 1 shows correlations between study variables. The observational estimate of the relationship between maternal educational attainment and child GCSE was 0.398 (95% CI: 0.368, 0.427). Using the best fitting polygenic score for years of education, the effect explained by genetic confounding was estimated at 0.073 (0.067, 0.080), corresponding to 18.2% of the total effect. After taking this genetic confounding effect (as captured by the polygenic score) into account, the relationship between maternal education and child GCSE was reduced to 0.324 (0.291, 0.357).

The sensitivity analysis is represented in Figure 1, where standardized estimates of the effect of maternal education on child GCSE are plotted as a function of the variance explained in the latter. We first re-estimated the effect of maternal education on child GCSE by adjusting for observed polygenic scores at different p value thresholds, explaining different amounts of variance in GCSE scores. We then estimated the effect of maternal education on child GCSE under scenarios in which polygenic scores could capture additional variance in educational outcomes (see Methods). The SNP-heritability scenario is based on the SNP-heritability of GCSE scores, which was previously estimated in TEDS to be 31% [10]. Under this scenario the effect of maternal education on child achievement further decreased to 0.175 (0.129, 0.222). The effect estimate was null under the twin-heritability scenario.

**Figure 1.**
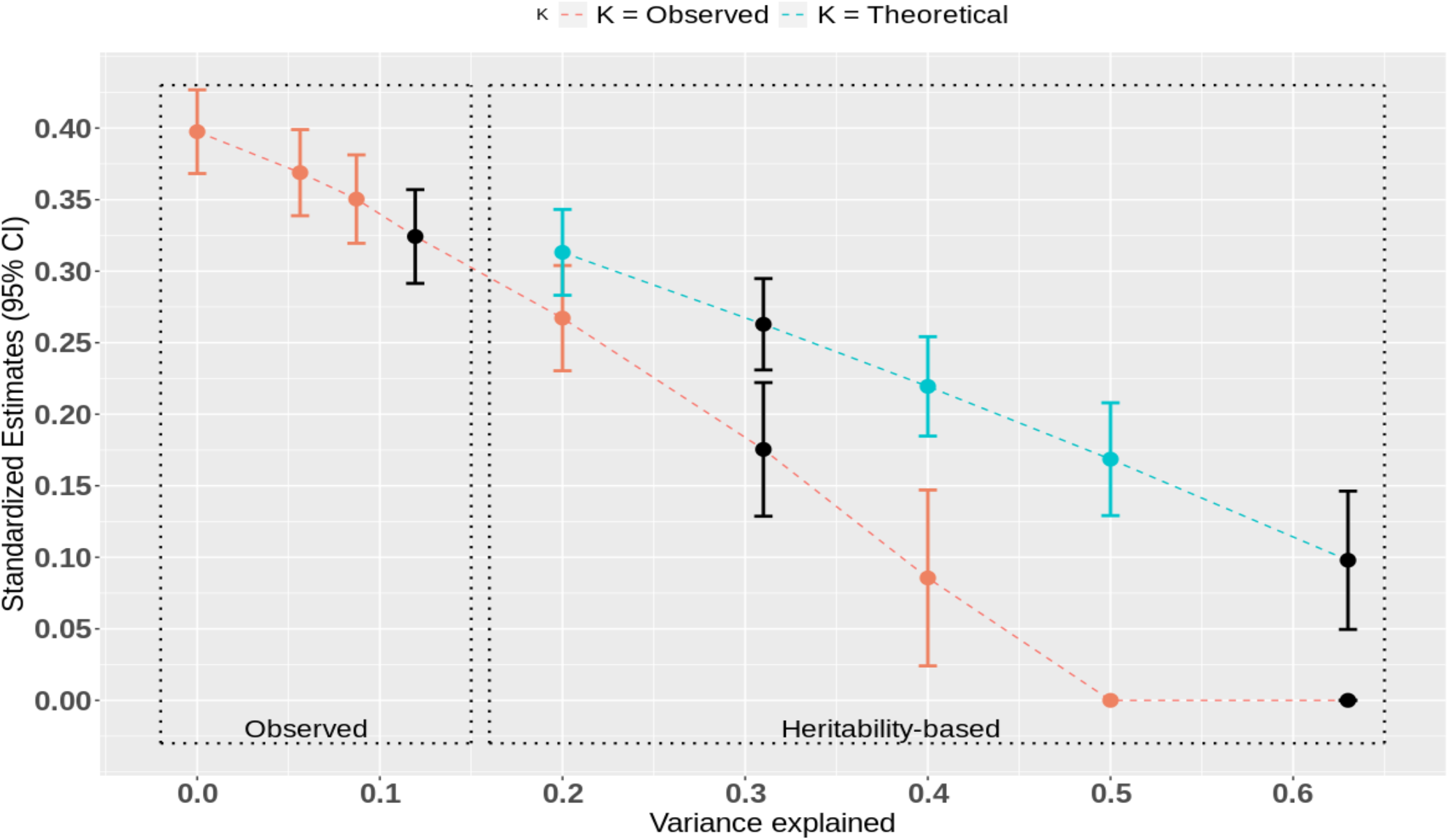
Gsens analysis of the effect of maternal educational attainment on child educational achievement Caption. Estimated standardized effect of maternal education on child educational achievement (Y axis) after accounting for genetic confounding using observed polygenic scores and heritability-based scenarios explaining an increasing percentage of variance (X axis). Point estimates and confidence intervals in black represent main estimates of interest, after accounting for (from left to right): 1: the best-fitting polygenic score; 2: SNP-heritability of educational achievement as assessed by GCSE scores in TEDS; 3: twin-heritability of educational achievement. A lower bound of 0 was imposed on the estimate, which is reached for the twin estimate of heritability (63%). The line “*k* = Observed” corresponds to heritability-based scenarios using values of model parameter *k* derived from observed polygenic scores (see Methods). “*k* = theoretical” corresponds to the value of *k* when the same trait is in parents and children and the heritability is the same in parents and children. In this case *k*=0.5 (see Methods).

We define *k* as the ratio of the standardized path from the polygenic score to the exposure divided by the standardized path from the polygenic score to the outcome (see Methods). The estimated k was 0.84. This is higher than the value of 0.5 expected when X and Y are the same trait measured in parents and children (meaning that the standardized association between the child polygenic score and maternal education should be, at most, half of the standardized association between the child polygenic score and the child educational outcome). In addition to sample-specific findings, this could be because the polygenic score for child educational achievement was derived from a GWAS of years of education in adults, which is closer to the maternal education phenotype (X) than the child GCSE phenotype (Y). A similar finding was observed by Bates et al. [20]. When setting *k* = 0.5 under the twin-heritability scenario, the estimate of 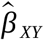 is still considerably reduced compared to the observed correlation but remains positive at 0.098 (0.066, 0.129).

#### Two polygenic scores: BMI and ADHD

The observational estimate of the relationship between maternal education and child BMI was *β_XY_* = −0.089 (−0.122, −0.057). Using the best fitting polygenic scores for years of education and BMI, the genetic confounding effect was estimated at −0.021 (−0.028, −0.013), corresponding to 23.0% of the total effect. After taking this genetic confounding effect into account, the relationship between maternal education and child BMI was −0.069 (−0.100, −0.037). The first scenario used SNP-based heritability estimates for years of education and BMI (see Table 1). In that scenario, the relationship between maternal education and child BMI further attenuated to −0.043 (−0.077,-0.009). In the twin heritability scenario, the estimate was null, meaning that, under this scenario, the entire association between maternal education and child BMI is accounted for by genetic confounding. Table 2 presents sensitivity analyses for BMI adjusting for both polygenic scores for the exposure and the outcome (GsensXY), only the outcome (GsensY), or only the exposure (GsensX). Estimates in bold are estimates from GsensXY reported in the text; other results presented in Table 2 are further explained in the next sections.

**Table 2.** Sensitivity analysis for BMI.

|  |  |  | GsensXY | GsensY | GsensX |
| --- | --- | --- | --- | --- | --- |
| Best Fitting PS | Unconstrained | Residual<sup>1</sup> | -0.081 (-0.114;-0.048) | -0.069 (-0.101;-0.037) | -0.089 (-0.123;-0.055) |
|  |  | G confound<sup>2</sup> | -0.009 (-0.021;0.004) | -0.021 (-0.029;-0.013) | 0.000 (-0.010;0.010) |
|  | Constrained | Residual | **-0.069 (-0.100;-0.037)** | -0.069 (-0.100;-0.037) | -0.089 (-0.123;-0.055) |
|  |  | G confound | **-0.021 (-0.028;-0.013)** | -0.021 (-0.028;-0.013) | 0.000 (-0.010;0.010) |
| SNP heritability | Unconstrained | Residual | -0.060 (-0.097;-0.022) | -0.028 (-0.065;0.009) | -0.089 (-0.123;-0.056) |
|  |  | G confound | -0.030 (-0.057;-0.002) | -0.061 (-0.086;-0.037) | 0.000 (-0.009;0.009) |
|  | Constrained | Residual | **-0.043 (-0.077;-0.009)** | -0.028 (-0.065;0.009) | -0.089 (-0.123;-0.055) |
|  |  | G confound | **-0.052 (-0.072;-0.033)** | -0.061 (-0.086;-0.037) | 0.000 (-0.010;0.009) |
| Twin heritability | Unconstrained | Residual | - | 0.132 (0.036;0.230) | -0.089 (-0.127;-0.051) |
|  |  | G confound | - | -0.222 (-0.320;-0.124) | -0.001 (-0.020;0.019) |
|  | Constrained | Residual | **0.00 (0;0)** | 0 (0;0) | -0.089 (-0.127;-0.051) |
|  |  | G confound | **-0.099 (-0.130;-0.067)** | -0.098 (-0.130;-0.065) | -0.001 (-0.020;0.019) |

*Note.* Estimates and 95% CI are provided for each scenario. <sup>1</sup>Residual association after adjusting for genetic confounding. <sup>2</sup>Genetic Confounding.

The observational estimate of the relationship between maternal education and child ADHD was-0.124 (−0.152, −0.096). Using the best fitting polygenic scores for years of education and ADHD, the genetic confounding effect was estimated to be −0.018 (−0.027, −0.009), corresponding to 14.3% of the total effect. After taking genetic confounding as captured by the polygenic scores into account, the relationship between maternal education and child ADHD attenuated to −0.106 (−0.135; −0.076). Table 3 shows results under different scenarios. In heritability-based scenarios, the relationship between maternal education and child ADHD was further attenuated, reducing to null in the twin-based scenario.

**Table 3.** Sensitivity analysis for ADHD.

|  |  |  | GsensXY | GsensY | GsensX |
| --- | --- | --- | --- | --- | --- |
| Best Fitting PS | Unconstrained | Residual<sup>1</sup> | -0.106 (-0.135;-0.076) | -0.117 (-0.145;-0.089) | -0.107 (-0.137;-0.077) |
|  |  | G confound<sup>2</sup> | -0.018 (-0.027;-0.009) | -0.007 (-0.010;-0.004) | -0.017 (-0.026;-0.008) |
|  | Constrained | Residual | **-0.106 (-0.135;-0.076)** | -0.117 (-0.145;-0.089) | -0.107 (-0.136;-0.077) |
|  |  | G confound | **-0.018 (-0.027;-0.009)** | -0.007 (-0.010;-0.004) | -0.017 (-0.026;-0.008) |
| SNP heritability | Unconstrained | Residual | -0.063 (-0.158;0.031) | -0.01 (-0.074;0.054) | -0.108 (-0.138;-0.079) |
|  |  | G confound | -0.061 (-0.153;0.032) | -0.114 (-0.173;-0.055) | -0.016 (-0.024;-0.008) |
|  | Constrained | Residual | **-0.052 (-0.096;-0.008)** | -0.01 (-0.074;0.054) | -0.107 (-0.136;-0.077) |
|  |  | G confound | **-0.078 (-0.113;-0.043)** | -0.114 (-0.173;-0.055) | -0.016 (-0.025;-0.008) |
| Twin heritability | Unconstrained | Residual | - | - | -0.089 (-0.123;-0.055) |
|  |  | G confound | - | - | -0.035 (-0.053;-0.017) |
|  | Constrained | Residual | **0 (0;0)** | 0 (0;0) | -0.089 (-0.123;-0.055) |
|  |  | G confound | **-0.129 (-0.158;-0.1)** | -0.127 (-0.156;-0.099) | -0.035 (-0.053;-0.017) |

*Note.* Estimates and 95% CI are provided for each scenario. <sup>1</sup>Residual association after adjusting for genetic confounding. <sup>2</sup>Genetic Confounding.

#### Model constraints

Tables 2 and 3 present constrained and unconstrained models. Unconstrained models, while closely fitting the data, can yield manifestly impossible values such as heritabilities above 100%, standardized paths above 1, or negative variances, in which case estimates are unreliable (which was the case for empty cells in Table 2 and 3). Implausible values are also observed. For example, in Table 2, the unconstrained GsensY estimate in the twin heritability scenario is positive, which would correspond to higher maternal education being linked to higher rather than lower BMI in children. This is understandable given that the best fitting polygenic score, which explains only 6.3% of variance in child BMI already explains 23% of the negative association between maternal education and child BMI. Under the twin heritability scenario, with BMI heritability of 64%, the association is likely to flip to a positive sign. This may be due to sampling. For example, if the cross path from BMI polygenic score to maternal education is, by chance, overestimated in TEDS, this will overestimate genetic confounding as captured by the polygenic score and impact estimation under the twin heritability scenario. It may also be because the true heritability of BMI is overestimated by the twin design. The constrained models therefore impose constraints on parameters to avoid impossible values (standardized paths above one and negative variances) and implausible values (cross-paths and residual associations flipping sign). While these models fit the observed data less well than the unconstrained models, they accommodate our prior beliefs about the plausible range of parameter values.

#### Bias amplification and simulations

*Gsens* uses estimates of heritability that provide useful benchmarks to estimate the extent of genetic confounding. It provides estimates of the strength of genetic confounding and of the residual association comprising the causal effect plus association through non-genetic confounders. However, these estimates are biased by the association between genetic and non-genetic confounders induced by conditioning on the exposure *X*, an instance of collider bias (see Methods). The bias is most pronounced when adjusting for polygenic scores that explain more variation in *X* than in *Y*, and may lead to estimates of residual association that are more biased than the observational association, a phenomenon termed bias amplification [21]. We therefore expect more bias amplification for GsensX compared to GsensY.

We conducted two sets of simulations based on the underlying causal model presented in Methods. In the first, we simulated polygenic scores capturing all genetic influences to examine the effect of bias amplification independent of measurement error. We compared estimates from the three *Gsens* functions to the true residual association net of genetic confounding (comprising the causal effect and non-genetic confounding). As shown in Figure 2, bias amplification can be particularly severe when there is no genetic confounding (top left panel, Figure 2), and when the heritability of X is high (bottom left panel, Figure 2). In such cases, adjusted estimates can indeed be higher than the observational association. In the presence of genetic confounding, the adjusted estimates are always closer to the true residual association than to the observational association (top and bottom right panels, Figure 2). However, bias is still present. Bias is greatest for GsensXY and GsensX. However, in all cases, estimates from GsensY have little or no bias, even when the heritability of X and the genetic correlation are high (bottom right panel, Figure 2). When there is no non-genetic confounding (0 on the X-axis), bias amplification does not occur and all *Gsens* estimators recover the causal effect.

**Figure 2.**
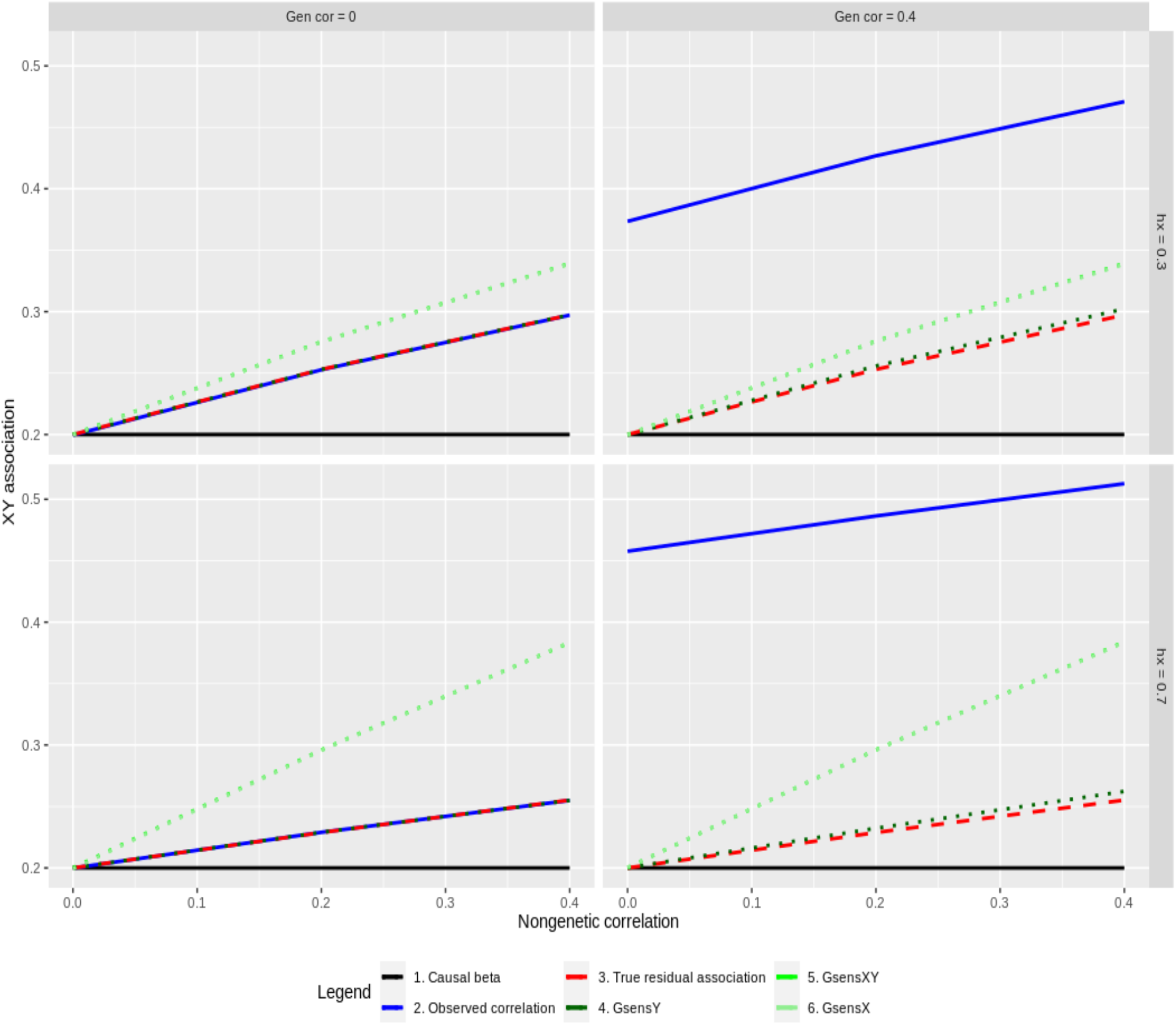
Bias amplification: simulation results. The standardized association between X and Y (Y-axis) is plotted as a function of the correlation between the non-genetic factors for X and Y that generate non-genetic confounding (X-axis). The figure is faceted left to right according to the genetic correlation that generates genetic confounding (gen cor), and top to bottom according to the heritability of X (hx). A subset of results is plotted with causal effect = 0.20, N = 10,000, heritability of Y = 70% and the full set of results is provided in S2. Note that estimates from GsensX and GsensXY are very similar in this first set of simulations but can differ in the second when polygenic scores are set to capture varying levels of the heritability of X and Y.

In the second set, we simulated polygenic scores that captured only a fraction of heritability. We simulated scenarios under which the polygenic score for X captures more of the variance of Y than the polygenic score for Y itself. This may happen with differentially powered GWAS. In our empirical example, the polygenic score for education captures almost as much variance in ADHD than the polygenic score for ADHD itself. Conceivably under such scenarios, using GsensX or GsensXY may better account for genetic confounding than GsensY. In our example, both GsensX and GsensXY using the polygenic score for education found a larger genetic confounding effect than when using the polygenic score for ADHD in GsensY alone. However, our simulations showed that, even under such circumstances, bias amplification under twin heritability scenarios remains larger for GsensX and GsensXY. Results are reported in S2.

In the two sets of simulations, constraints imposed on the model to avoid impossible or implausible results often removed part or all bias. However, we caution against systematically applying constraints as they do not necessarily remove bias and may artificially reduce standard errors. Results of constrained models, as well as standard errors for all models, are reported in S2. Of note, standard errors for GsensY were systematically lower than for GsensX and GsensXY. Empirical standard errors closely matched the analytic estimates.

These results suggest that GsensY should be preferred in all situations and that *Gsens* is best adapted when the outcome of interest is strongly predicted by its polygenic score. When the polygenic score for the exposure is more predictive, GsensXY can be used, particularly to estimate genetic confounding with observed polygenic scores, but should be interpreted with caution under heritability scenarios. Importantly, simulations show that small remaining bias for GsensY is conservative in the sense that it underestimates genetic confounding. This lessens the risk of overadjusting the association between X and Y.

## Discussion

In the present study, we combined polygenic scores with heritability estimates in a genetic sensitivity analysis – *Gsens –* aiming to gauge to what extent genetic confounding can account for observed epidemiological associations. The genetic sensitivity analysis we propose adds a new tool to probe the robustness of associations between exposures and outcomes. This approach requires a genotyped cohort with relevant phenotypic measurements, which is increasingly the rule for epidemiological cohorts rather than the exception. It is therefore possible to envisage *Gsens* analysis becoming routine. Below, we first discuss our empirical findings regarding the associations between maternal education and child educational attainment, BMI, and ADHD. We then discuss the interpretation and applicability of *Gsens.*

### Maternal education and developmental outcomes

Our findings show that the association between maternal education and developmental child outcomes were still present under a SNP-heritability scenario but were null under a twin-heritability scenario. Overall, the observed association between maternal education and these three developmental outcomes may largely be due to genetic confounding.

Relevant to our findings is previous research using causal inference designs to investigate the effect of parental educational attainment on child educational attainment. In particular, a systematic review on the topic has summarized evidence from twin and adoption designs, as well as non-genetic instrumental variable estimations [22]. The systematic review suggested that intergenerational associations between parent and child educational attainment are largely driven by selection effects, including genetic confounding; it suggests only small but still significant causal effects. A new method – the ‘virtual-parent design’ – has recently emerged, which consists of splitting parental genetic variants associated with a parental exposure into variants transmitted and nontransmitted to the child [20,23]. Parental polygenic scores including only nontransmitted variants are free from genetic confounding and index plausible causal effects of the parental exposure on the child outcome. Empirical findings implementing this method in education research suggest substantial genetic confounding and small but still significant causal effects of parental attainment on child attainment. Our findings on educational attainment are consistent overall with this literature. Although genetic confounding accounted entirely for the association between parental education attainment and child achievement, we detected a small but significant effect when using the upper theoretical limit of the *k* parameter, consistent with previous findings. In addition, scenarios based on lower heritability estimates also yielded small but significant effects, raising the possibility that null findings resulted from overestimated twin heritability estimates, a possibility further discussed below. Taken together, this set of findings represents a clear call for caution when interpreting non-genetically informed epidemiological studies on the role of maternal education.

### Interpreting the sensitivity to genetic confounding analysis

Two key points regarding the interpretation of *Gsens* findings must be highlighted. First, the reliability of findings depends on the accuracy of heritability estimates. Overestimating heritability would lead to overestimating genetic confounding and thus underestimating the residual association between exposure and outcome. For example, a recent study of 193,518 twins across 16 countries has showed that educational attainment is 43% heritable [24]. As can be seen in Figure 1, this lower estimate would lead to a substantially larger adjusted association. Improved estimates of heritability that better reflect true genetic effects can be plugged into our method as they become available. In addition, power increase in GWAS will improve the reliability of *Gsens* in the following ways: (i) by increasing the predictive power of polygenic scores and therefore the accuracy of observed scenarios; (ii) by improving the accuracy of parameter estimates for the sensitivity analysis.

Second, *Gsens* aims to estimate genetic confounding and the residual association net of genetic confounding. This is distinct from the genetic overlap as estimated by bivariate twin or mixed-model methods, which decomposes the phenotypic association into genetic and environmental components (the percentage of the phenotypic correlation that is due to genetic overlap is called bivariate heritability). This is because a genetic correlation can arise from genetic effects on the outcome mediated via the exposure and the causal path (i.e. mediated pleiotropy) or by direct genetic effects on the outcome (i.e. unmediated pleiotropy). In twin-based methods, the corresponding residual association is an environmental association in that it excludes all genetic components from the observed association, including those mediated by the causal effect. In contrast, *Gsens* aims to remove only the genetic confounding or unmediated pleiotropy. In the absence of nongenetic confounding, the *Gsens* residual association would correspond to the causal effect, while the environmental association would be lower than the causal effect. Further clarifications regarding these conceptual differences can be found in S1 Annex 1.

In contrast, a conceptually similar approach to ours is taken in the latent causal variable (LCV) model [25], which also estimates genetic confounding parameters without identifying the causal effect of the exposure. The emphasis of LCV is however on comparing the genetic confounder effects on the risk factor to those on the outcome. Mendelian Randomization (MR) methods impose stronger assumptions on the genetic confounding effects to explicitly identify the causal effects [26]. In comparison with MR and LCV methods, our approach requires only the standard assumptions of structural equation modelling, and retains much of the precision of the observational association. However we cannot identify the causal effect from the residual association, unless we assume no non-genetic confounding. We contend that where there is substantial genetic confounding, our approach provides insights into the likely existence and magnitude of a causal effect. Indeed, our approach recreates a standard regression adjustment for a polygenic score explaining the entire heritability, should such a score be available. In this respect we follow traditional lines of sensitivity analysis in epidemiological studies, accepting that bias may still remain from residual confounding.

### Limitations and research avenues

As TEDS does not include parental genotypes, we did not model their role directly. Where such data are available, *Gsens* could be extended in the intergenerational context. However, although our examples were of intergenerational associations, Gsens can be generally implemented when both the exposure and the outcome are measured for the same individuals, without considering parental effects. Furthermore, providing the sampling populations are exchangeable, it is not necessary for the genetic and observational associations, nor the heritabilities, to be estimated in the same data sets. *Gsens* may be extended in future to explicitly consider measured confounders, which may further reduce residual amplification bias due to nongenetic confounding.

### Conclusions

We propose a genetic sensitivity analysis aiming to adjust for genetic confounding in epidemiological associations. Gsens implements structural equation models to adjust for genetic confounding based on polygenic scores and heritability estimates. Gsens can be applied as long as a suitable GWAS for the outcome is available, even when a GWAS for the exposure is not available or does not provide adequate instruments for MR analyses. For example, Gsens can be applied to test whether the association between urbanicity and schizophrenia is susceptible to genetic confounding based on the polygenic score and heritability estimates for schizophrenia. We therefore propose that Gsens can be conceived as a complementary method, suited for complex environmental exposures that are of interest for health, behavioural and social sciences.

## Methods

### Participants

Participants were drawn from the Twins Early Development Study (TEDS), a longitudinal study of twin pairs born in England and Wales, between 1994 and 1996. Detail regarding TEDS, the recruitment process, and representativeness can be found elsewhere [27]. A total of 7,026 unrelated individuals have been genotyped in TEDS. For each individual analysis, sample size comprised between 3,663 to 4,693 individuals with data for maternal education and each outcome. Written consent was obtained from all the families who agreed to take part in the study. This study was approved by the Institute of Psychiatry, Kings College London, Ethics Committee.

### Measures

Maternal educational attainment was reported by mothers at first contact, when children were on average 18 months old, with 8 levels: 1= no qualifications; 2 = CSE grade 2-5 or O-level/GCSE grade D-G; 3 = CSE grade 1 or O-level/GCSE grade A-C; 4 = A-level or S-level; 5 = HNC; 6 = HND; 7= undergraduate degree; 8 = postgraduate qualification.

*Child educational achievement* was operationalized as performance on the standardized UK-wide examination, the General Certificate of Secondary Education (GCSE), at 16 years. We computed a mean of the three compulsory core subjects, mathematics, English, and science (further details in [10]). A total of 3,785 genotyped individuals had data on both maternal education and child GCSE.

Body Mass Index (BMI) was derived from parent reported (ages 11 and 14 years) and self-reported weight and height (age 16 years). Extreme BMI values (<1% and >99% quantiles) were winsorized and resulting values were averaged across ages 11 to 16 years. A total of 3,663 genotyped individuals had data on maternal education and the resulting BMI score.

The DSM-IV ADHD symptom subscale, taken from the Conners’ Parent Rating Scales–Revised [28] was completed by mothers to assess inattentive and hyperactive/impulsive symptoms (9 for hyperactivity/impulsivity and 9 for inattention). Each item was rated on a 4-point Likert scale ranging from 0 (not at all true) to 3 (very much true). A total ADHD score was created by averaging scores across the following mean ages of participants at assessments: 8, 11, 14, and 16 years. The score measures population symptoms dimensionally and not the clinical disorder. A total of 4,693 genotyped individuals had data on maternal education and the ADHD score.

### Analyses

Genotyping, quality control procedures and principal component analysis are detailed in S1 section 1. A total sample of 7,026 participants with European ancestry remained after quality control. Single Nucleotide Polymorphisms (SNPs) were excluded if the minor allele frequency was <5%, if more than 1% of genotype data were missing, or if the Hardy Weinberg p-value was lower than 10^−5^. Non-autosomal markers and indels were removed.

We computed genome-wide polygenic scores based on summary statistics from the following genome-wide association studies (GWAS): (i) years of education [29]; (ii) ADHD [13]; and (iii) BMI [11]. In some cases, like ADHD, the GWAS phenotypes do not match our measures exactly; however they still explain substantial variation and can be extrapolated up to the heritability of our measure. Polygenic scores for all TEDS participants and all traits were computed using PRSice software [15], with prior clumping to remove SNPs in linkage disequilibrium (r^2^ > 0.10). PRSice allowed us to select the best-fitting polygenic score for each trait, e.g. maximizing the amount of variance explained by the polygenic score for BMI in TEDS participants' BMI. To this end, we computed a series of polygenic scores including an increasing number of SNPs corresponding to increasing p-value thresholds (e.g. all SNPs associated to BMI at p <.0001 and p <.10) as illustrated in S1 eFigures 1, 2, & 3. Using linear regression analyses, we estimated the proportion of variance explained by each generated polygenic score in the corresponding trait in TEDS. The following covariates were included in regression analyses: sex, age (for GCSE), and 10 principal components of ancestry.

**Figure 3.**
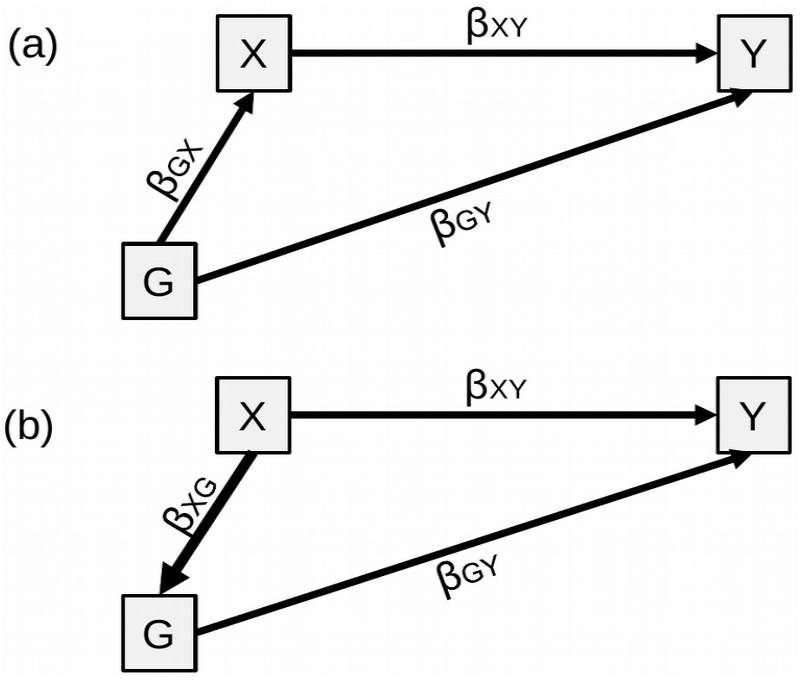
Genetic confounding, one polygenic score case. Caption. Figure 3 (a) represents the underlying model. (b) represents the model to calculate the confounding effect by treating G as a 'mediator’. Of note is that the commonly-used terminology ‘genetically mediated’ can be confusing. Although ‘genetically mediated’ makes sense statistically, conceptually, a mediator is on the causal pathway from the predictor to the outcome. However, because germline genetic variants are set at conception and do not change throughout the lifespan, posterior exposures (e.g. individual alcohol intake) cannot influence health outcomes (e.g. depression) through modifying germline genetic variants [32]. Although statistically treated as a mediator here to estimate confounding, conceptually G does not qualify as a mediator. Variances not represented for simplicity.

### Genetic confounding

We assume a linear structural equation model framework [30]. Akin to third variable confounding, genetic confounding is represented in Figure 3a: genetic factors (G) – here measured by polygenic score(s) – influence both the exposure (X) and the outcome (Y). MacKinnon et al. demonstrated that mediation and confounding are statistically identical in linear structural equation modelling [31].

Therefore, genetic confounding can be estimated by treating the confounder– here the polygenic score G as a mediator of the effect of X and Y (Figure 3b). The confounding effect is the indirect effect of X on Y through G: *β_XG_ β_GY_* . We also calculated the proportion of the observed effect of X on Y that is accounted for by genetic confounding, i.e.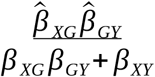

When the polygenic scores for the predictor (G1) is different from the polygenic score for the outcome (G2), the confounding effect is estimated in a similar fashion as the sum of all the indirect effects from X to Y through G1 and/or G2 (Figure 4a and 4b).

**Figure 4.**
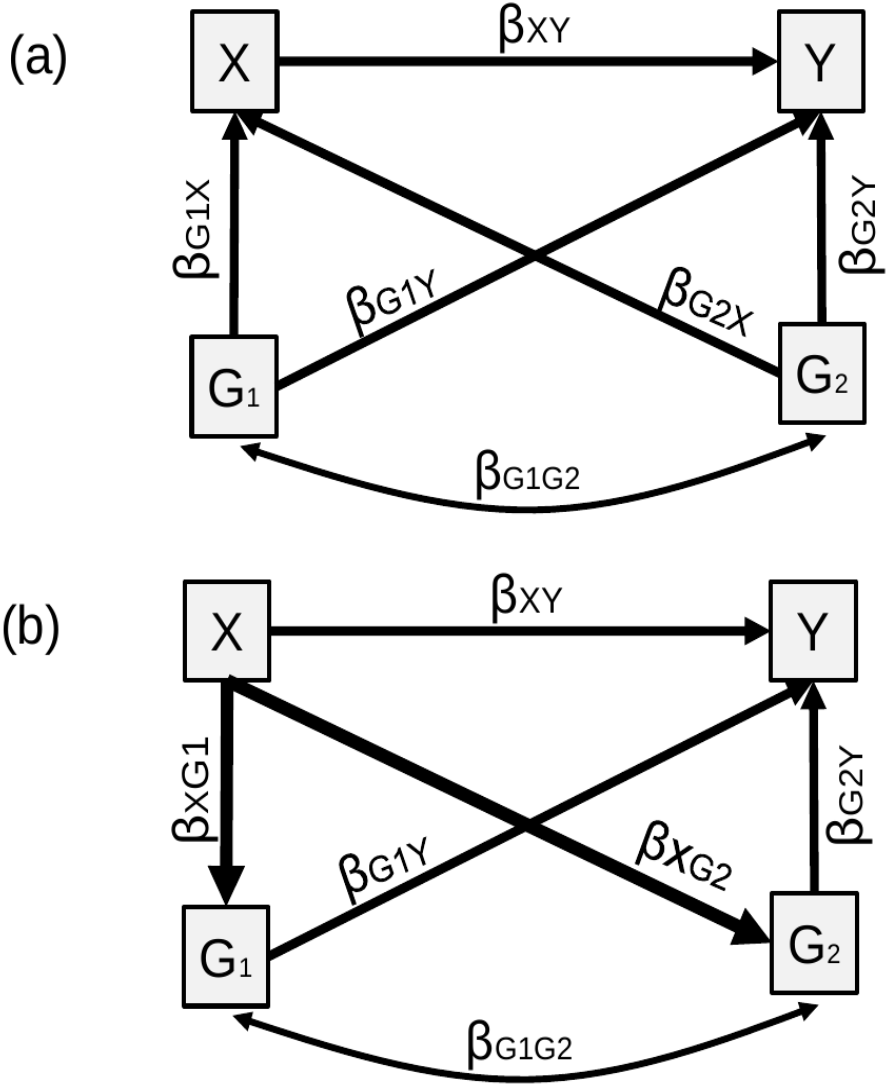
Genetic confounding, two polygenic score. Caption. Figure 4a represents the underlying causal model. Figure 4b represents the model to calculate the confounding effect, which is equal to: 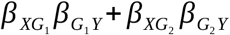. Note that when model variables are standardized, the genetic confounding effect can also be obtained based on 4a by 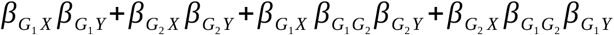. Variances not represented for simplicity.

**Figure 5.**
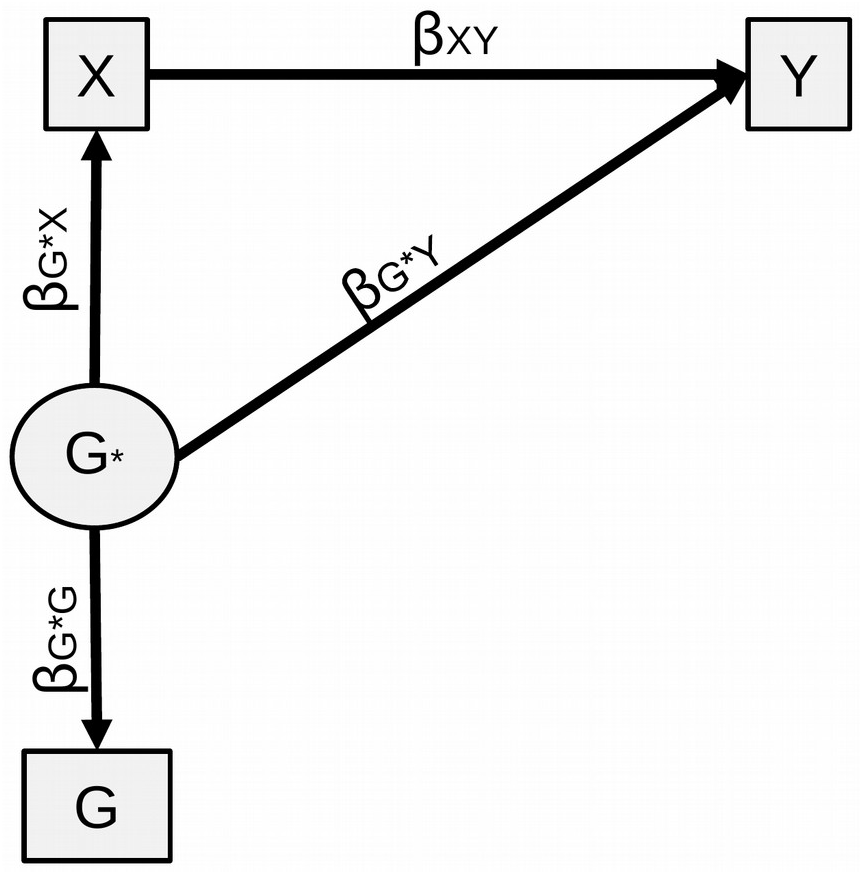
Sensitivity analysis, one polygenic score. When using the best-fitting polygenic score, *β_XY_* can be estimated using standardized associations between the observed polygenic score, X and Y, as in expression (1). In the sensitivity analysis, a structural equation model is fitted with a latent variable G* representing heritability, as presented in Figure 5. This can be understood as correcting for measurement error, i.e. G being an imperfect measure of G*. Genetic confounding estimated under this model reflects heritability under the chosen scenario rather than only what is captured by the polygenic score. We fitted structural equation models using the R package ‘lavaan’ [33]. The latent variable is set to capture the heritability of Y under the sensitivity analysis scenario (e.g. twin-heritability). The following constraint is is applied: 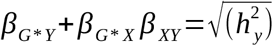Variances not represented for simplicity.

Genetic confounding effects were calculated for all three developmental outcomes:

- Maternal education to child educational achievement using the best-fitting polygenic score for years of education (as in Figure 3);
- Maternal education to child BMI using best-fitting polygenic scores for years of education (G1) and BMI (G2) (as in Figure 4);
- Maternal education to child ADHD symptoms using best-fitting polygenic scores for years of education (G1) and ADHD symptoms (G2) (as in Figure 4).

In these analyses, the effect size of X on Y decreases as a function of the strength of genetic confounding. However, this approach does not account for all the genetic confounding. This is because polygenic scores based on current GWAS capture a relatively small amount of all genetic influences. For example, the current polygenic score for BMI explains around 6% of the variance in BMI in TEDS, far less than SNP-based and twin heritability estimates of BMI heritability. The sensitivity analysis we propose aims to address this issue.

### Sensitivity analysis

The sensitivity analysis aims to answer the following question: is X is associated with Y after we control for all genetic confounding? In other words, to what extent would *β_XY_* decrease if we were to control for ‘perfect’ polygenic scores capturing all genetic influences on X and Y rather than a small fraction. This is done by estimating *β_XY_* under plausible scenarios that combine information on existing polygenic scores and heritability estimates.

#### Single polygenic score

For maternal education and child educational achievement, we used a polygenic score for the child, derived from the GWAS of years of education, which predicts a substantial amount of variance both in child GCSE but also in maternal education. The effect of maternal education on child educational attainment can be first adjusted for the observed best-fitting polygenic score. However, this polygenic score does not capture all the heritability of the outcome and therefore incompletely adjusts for genetic confounding. The sensitivity analysis consists in re-examining the effect of maternal education under scenarios where the polygenic score could capture additional variance in child GCSE up to SNP-based and twin-based estimates of heritability.

Figure 3a shows the underlying model of relationships between the polygenic score (G), the predictor (X) and the outcome (Y). We can obtain an adjusted effect of X on Y based on the observed associations available with the following expression:

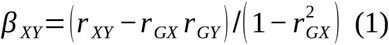

where *β_XY_* is the adjusted effect and *r* denotes observed standardized associations. Details are presented in S1 section 2. Importantly, *β_XY_* corresponds to the standardized association between X and Y minus genetic confounding, i.e. the residual association between X and Y net of genetic confounding. In other words, *Gsens* removes only genetic confounding and not all genetic effects shared between X and Y, which comprise both genetic confounding and genetic effects on Y mediated by X via a causal pathway. When subtracting all shared genetic effects, including those arising from the causal effect, the residual association becomes the 'environmental association'. This is similar to what happens in bivariate decomposition of the phenotypic correlation in twin and mixed model designs and is distinct from *Gsens* estimates. This distinction is further clarified in S1 Annex 1.

#### Complete genetic confounding

In equation (1), the association between X and Y is completely genetically confounded when the adjusted effect *β_XY_* = 0. We can then express the observed standardized association as a function of the heritabilities of X and Y under complete genetic confounding as:

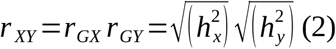

When the adjusted effect of X on Y is null, then *r XY* is equal to genetic effects through G. In the special case where X and Y are the same trait in parent and child and assuming constant heritability across generations, we thus obtain:

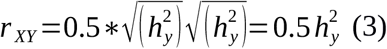

This means that the adjusted effect of X on Y is likely to be null whenever the observed association does not exceed half of the trait heritability. As such, a given association between parental and child traits can be assessed against Figure 6 and if it lies in the shaded area, it is consistent with complete genetic confounding. Importantly, associations not in the shaded area can still be confounded by environmental exposures. See S1 section 3 for additional details on equations (2) and (3).

**Figure 6.**
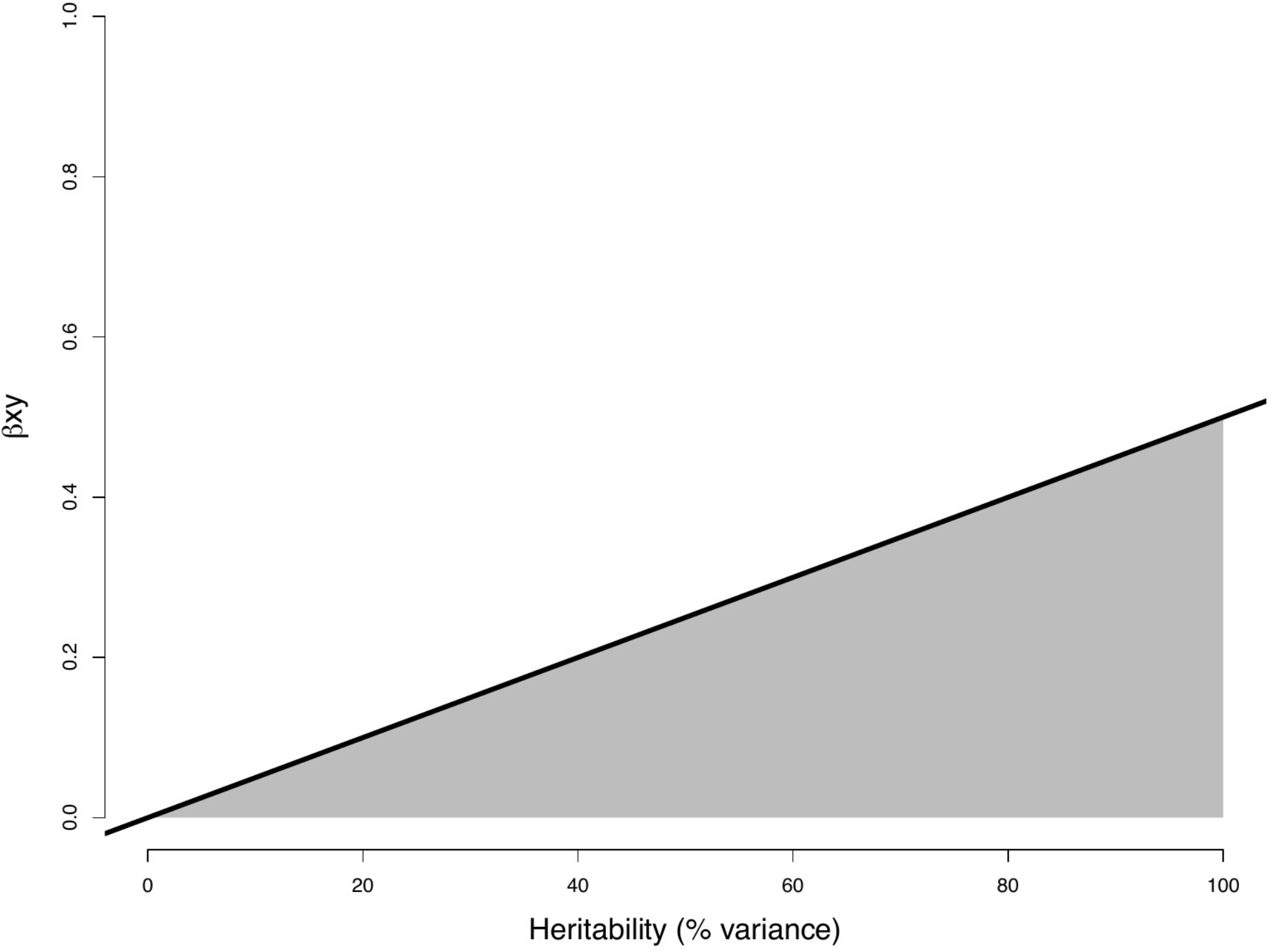
The role of genetics in explaining phenotypic associations between parent and child. *Caption.* Standardized observed associations between the same traits in the mother (or father) and the child are represented as a function of trait heritability. An observed association of 0.20 with trait heritability of 0.60 is consistent with complete genetic confounding. Conversely, an association of 0.40 with heritability of 0.40 is not consistent with complete genetic confounding .

### The two polygenic scores case

When predictor and outcome are different variables – for example maternal education and child BMI – two polygenic scores can be used in the sensitivity analysis, as shown in Figure 4. In theory, if we had a polygenic score capturing all genetic influences for Y, this score would also capture all the genetic overlap between X and Y, and we could use the one polygenic score case above. In practice, polygenic scores do not capture all genetic influences on their respective phenotypes and are differentially powered, which is why we examine the utility of a two polygenic scores solution. In the two polygenic scores case, new parameters are introduced including the cross paths from each polygenic score to the other phenotype (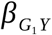 and 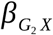). Due to these new parameters, the derivation of *β_XY_* becomes more complex than for the single case polygenic score. We thus generalize the structural equation model to two latent variables and polygenic scores as in Figure 7. Further details in S1 Section 4.

**Figure 7:**
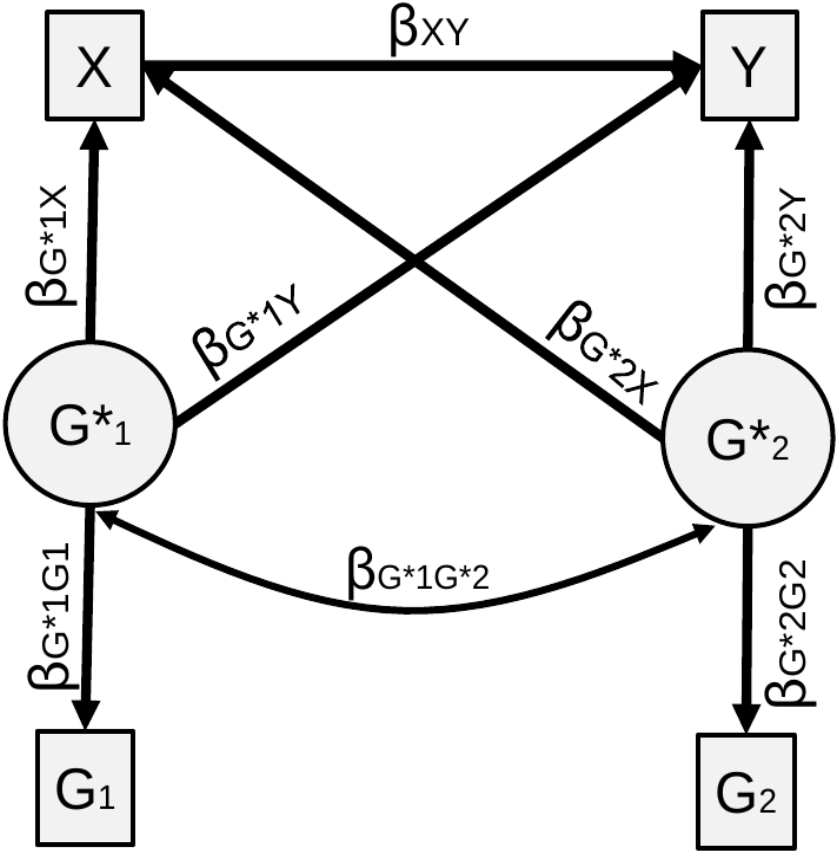
Sensitivity analysis, two polygenic scores. The following constraints are imposed on the model: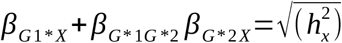 and 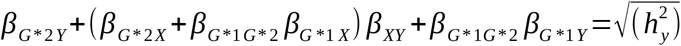 Variances not represented for simplicity.

### Model assumptions

Our approach requires the standard assumptions of structural equation modelling, including normality of the observed and latent variables and no unmodelled confounding or interaction effects. For polygenic traits the normality assumptions are reasonable. Note that although polygenic scores are constructed from additive models, we make no such assumption for the true latent genetic value, only that it has a linear relationship with the polygenic score. Unmodelled confounders can create bias amplification, as we show in our simulations. However note that all heritable confounders would be included in the latent genetic values under the heritability scenarios, and so only the non-genetic components of unmodelled confounders would create bias.

### Simulations

In order to verify the performance of *Gsens* under its own assumptions, and to study the possibility of bias amplification, we conducted simulations based on the underlying causal model presented in Figure 8. Simulations were conducted with the SimulateData() function in package 'lavaan' embedded within the wrapper simulation package SimDesign [34,35].

**Figure 8.**
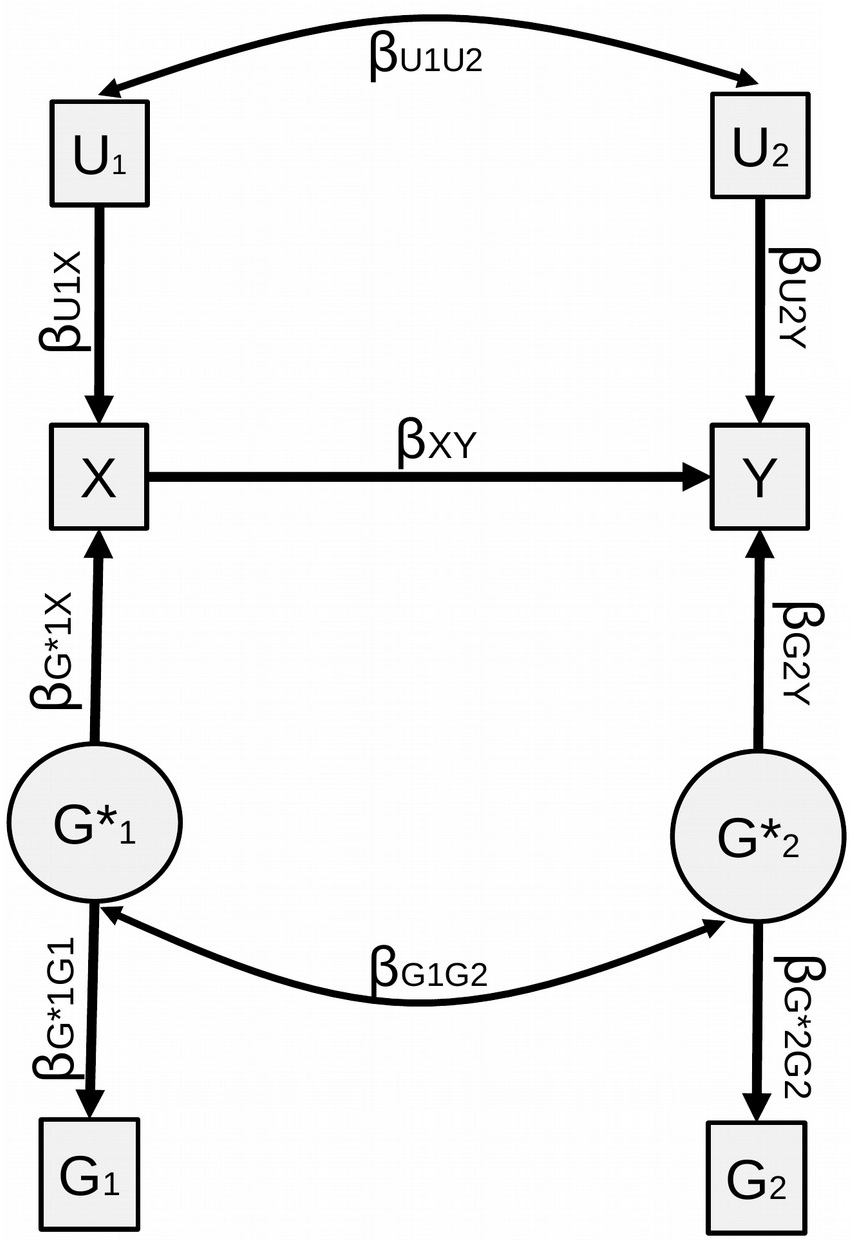
Simulation generative model. The figure represents the generative model for simulations.

In the first set of simulations, loadings from 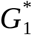 to *G* and 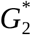 to *G* were fixed to unity (there by simulating polygenic scores capturing the whole heritability) in order to examine amplification bias independently of the latent structure of the model. We chose parameters based on reasonable values, with the following combinations: X and Y were 30% or 70% heritable, and influenced by respective non-genetic influences of 55% or 15% (leaving 15% of unexplained variance); genetic and non-genetic correlations of 0 or 0.40; a causal effect of 0 or 0.2.

In the second set of simulations, we fixed the causal effect to 0.20 and heritabilities to 70% but values of the loadings were set so that the resulting polygenic scores *G*_1_ and *G*_2_ would capture 1% or 10% of the variance of X and Y, respectively. This resulted in either polygenic scores with equal explanatory power or asymmetric situations where, e.g. one polygenic score explained 10% of the variance in X and the other polygenic score explained only 1% of the variance in Y. In this case, the resulting association between the first polygenic score and Y may actually be greater that the association between the second and Y, which can result in non-null cross-paths. Such a situation can stem from the differential accuracy of GWAS for X and Y.

## Acknowledgments

We gratefully acknowledge the ongoing contribution of the participants in the Twins Early Development Study (TEDS) and their families. This project has received funding from the European Research Council (ERC) under the European Union’s Horizon 2020 research and innovation programme (grant agreement No. 863981) and the MRC (MR/S037055/1). We are grateful to Dr. Jessie Baldwin for comments on the manuscript and the tutorial.

