## Supplement 1 for "Genetic sensitivity analysis: adjusting for genetic confounding in epidemiological associations"

#### Section 1. Genotyping protocol and quality control

DNA for 8,743 individuals (including 3,722 dizygotic co-twin samples) was extracted from saliva and buccal cheek swab samples and hybridized to HumanOmniExpressExome-8v1.2 genotyping arrays at the Institute of Psychiatry, Psychology and Neuroscience Genomics & Biomarker Core Facility. The raw image data from the array were normalized, pre-processed, and filtered in GenomeStudio according to Illumina Exome Chip SOP v1.4. (http://confluence.brc.iop.kcl.ac.uk:8090/display/PUB/Production+Version%3A+Illumina+Exome+Chip+SOP+v1.4). In addition, prior to genotype calling, 919 multi-mapping SNPs and 501 samples with callrate <0.95 were removed. The ZCALL program (see Web resources section) was used to augment the genotype calling for samples and SNPs that passed the initial QC.

DNA from 3,747 samples was extracted from buccal cheek swabs and genotyped at Affymetrix, Santa Clara, California, USA. From this sample, 3,665 samples were successfully hybridized to AffymetrixGeneChip 6.0 SNP genotyping arrays (http://www.affymetrix.com/support/technical/datasheets/genomewide_snp6_datasheet.pdf) using experimental protocols recommended by the manufacturer (Affymetrix Inc., Santa Clara, CA). The raw image data from the arrays were normalized and pre-processed at the Wellcome Trust Sanger Institute, Hinxton, UK for genotyping as part of the Wellcome Trust Case Control Consortium 2 (https://www.wtccc.org.uk/ccc2/) according to the manufacturer's guidelines (http://www.affymetrix.com/support/downloads/manuals/genomewidesnp6_manual.pdf). Genotypes for the Affymetrix arrays were called using CHIAMO (https://mathgen.stats.ox.ac.uk/genetics_software/chiamo/chiamo.html).

After initial quality control and genotype calling, the same quality control was performed on the samples genotyped on the Illumina and Affymetrix platforms separately using PLINK (Chang et al., 2015; Purcell et al., 2007), R (R Core Team, 2017), BCFtools (Li, 2011), and EIGENSOFT (Patterson, Price, & Reich, 2006).

Samples were removed from subsequent analyses on the basis of call rate (<0.98), suspected non-European ancestry, heterozygosity, and relatedness other than dizygotic twin status. SNPs were excluded if the minor allele frequency was smaller than 0.5%, if more than 2% of genotype data were missing, or if the Hardy Weinberg *p*-value was lower than $10^{-5}$. Non-autosomal markers and indels were removed. Association between SNP and the platform, batch, plate or well on which samples were genotyped was calculated; SNPs with an effect *p*-value < $10^{-4}$ were excluded. A total sample of 10,346 samples (including 3,320 dizygotic twin pairs and 7,026 unrelated individuals), with 7,289 individuals and 559,772 SNPs genotyped on Illumina and 3,057 individuals and 635,269 SNPs genotyped on Affymetrix remained after quality control.

Genotypes from the two platforms were separately phased using EAGLE2 (Loh et al., 2016), and imputed into the Haplotype Reference Consortium (release 1.1) using the Positional Burrows-Wheeler Transform method (Durbin, 2014) through the Sanger Imputation Service (McCarthy et al., 2015). Prior to merging, we excluded variants with info <0.75 and removed non-overlapping SNPs between platforms. After merging, we tested for minor allele frequency differences between platforms and removed SNPs with an effect p-value < $10^{-4}$, and Hardy Weinberg p-value > $10^{-5}$. Using these criteria, 7,363,646 genotyped and well-imputed SNPs were retained for the analyses.

We performed principal component analysis on a subset of 39,353 common (MAF > 5%), perfectly imputed (info = 1) autosomal SNPs, after stringent pruning to remove markers in linkage disequilibrium ($r^2 > 0.1$) and excluding high linkage disequilibrium genomic regions so as to ensure that only genome-wide effects were detected.

**eFigure 1: Predictive accuracy of the polygenic score for years of education predicting child GCSE in TEDS for different P-value thresholds**

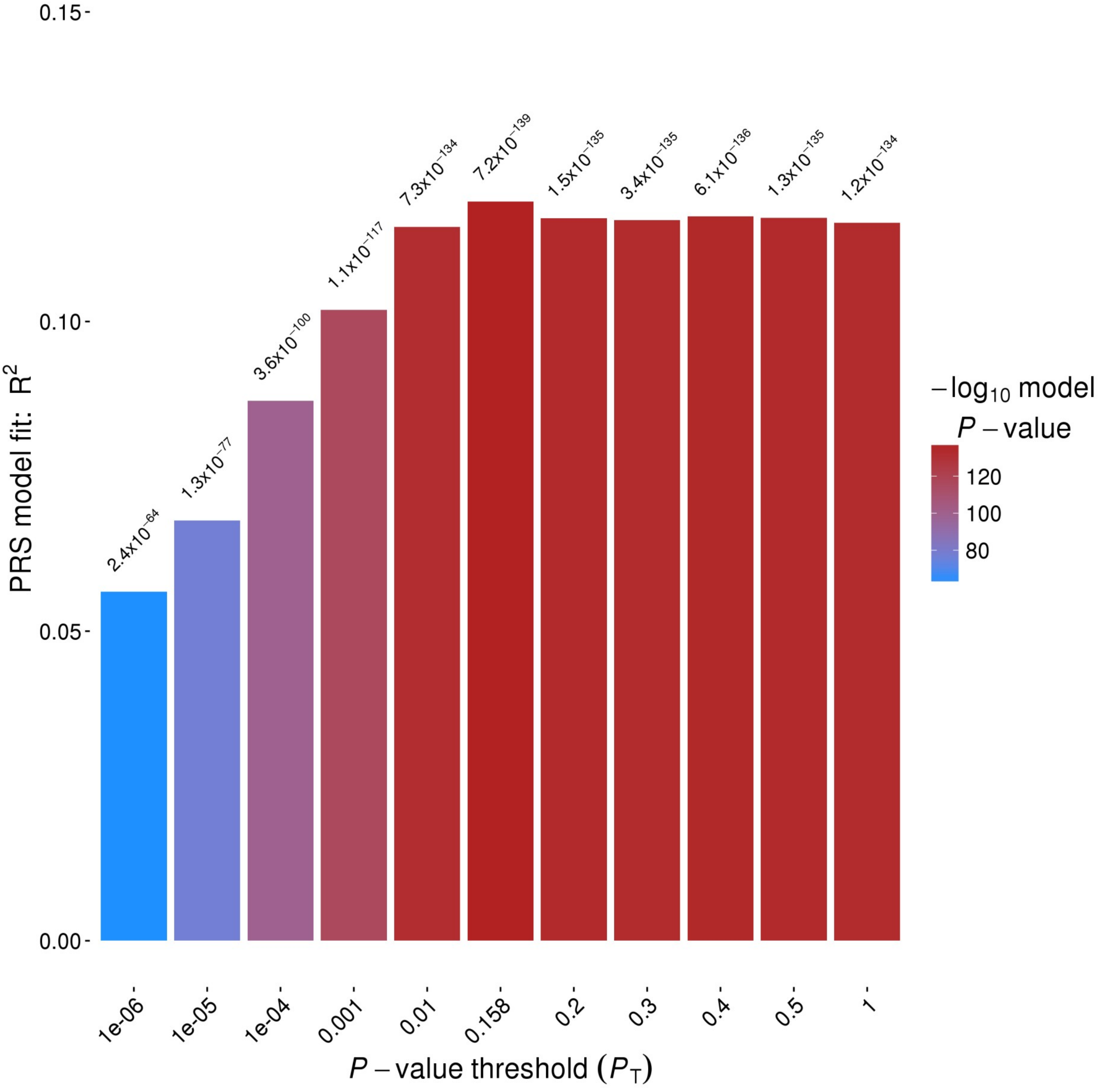

**eFigure 2: Predictive accuracy of the polygenic score for BMI predicting child BMI in TEDS for different P-value thresholds**

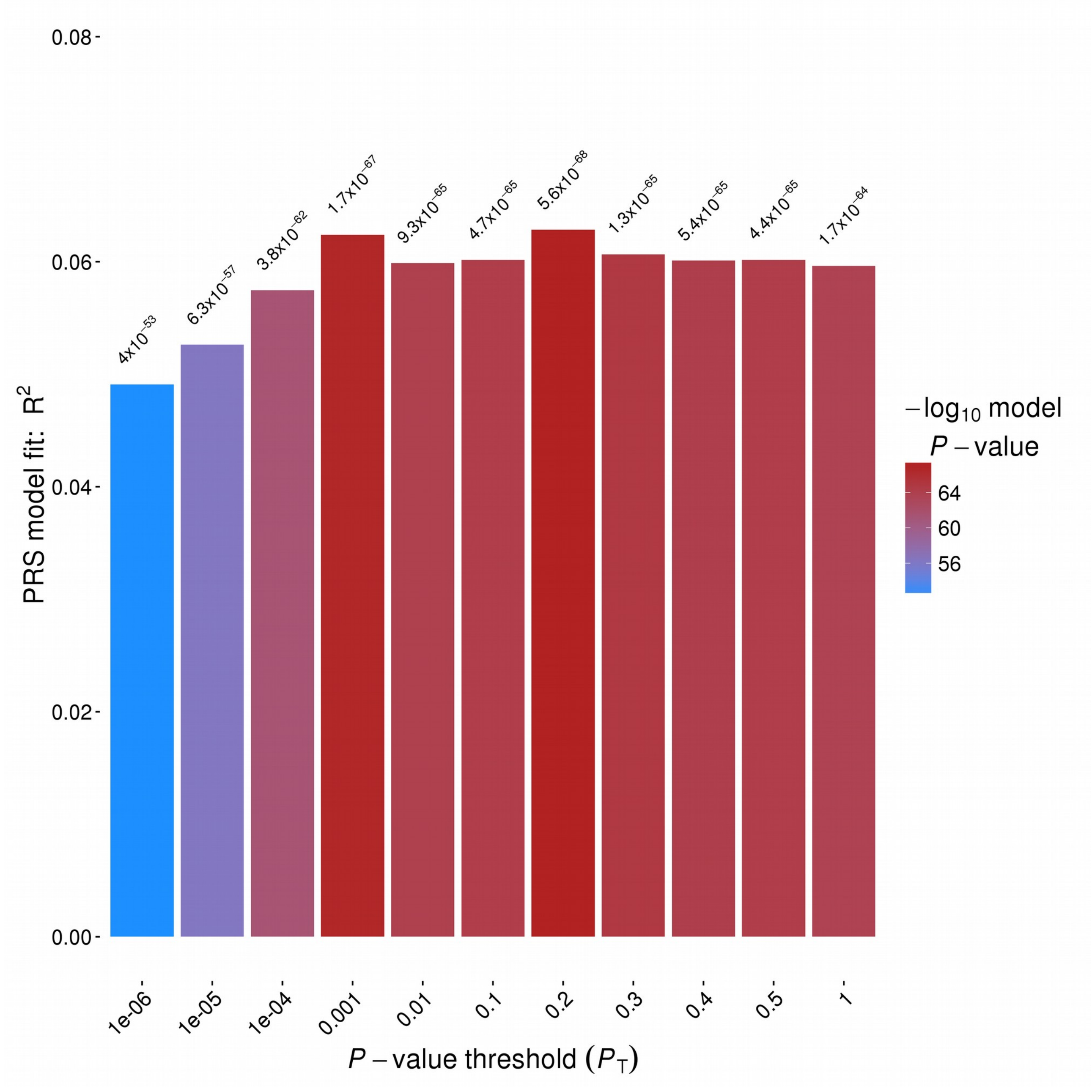

#### eFigure 3: Predictive accuracy of the polygenic score for binary ADHD predicting ADHD symptoms in TEDS for different P-value thresholds

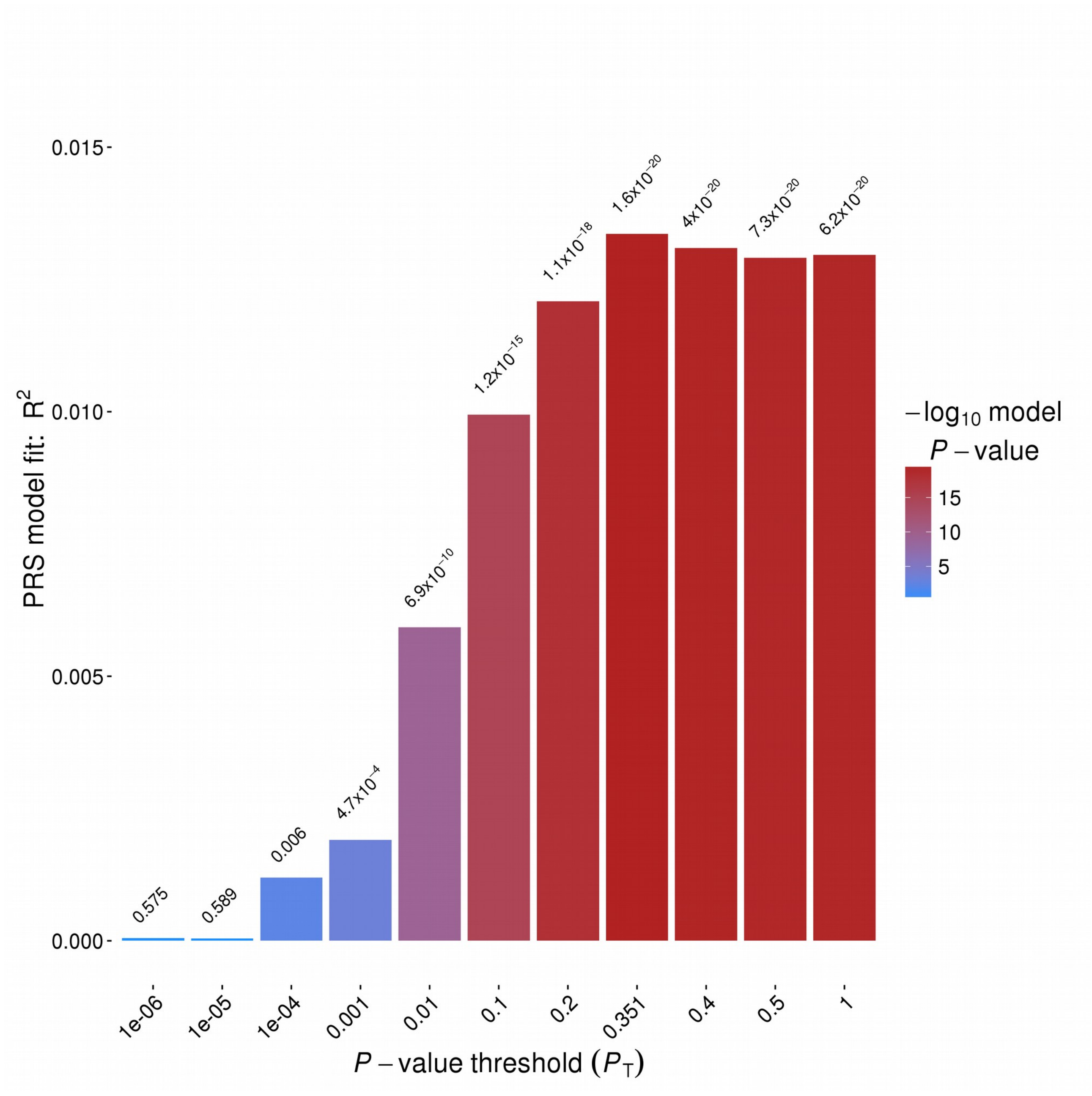

#### eTable 1. Correlations between study variables

| | Maternal Education | GCSE | BMI | ADHD | EDU PS | BMI PS | ADHD PS |
|---|---|---|---|---|---|---|---|
| Maternal Education | 1 | 0.3975 | -0.0894 | -0.1240 | 0.2909 | -0.0839 | -0.0630 |
| GCSE | 0.3975 | 1 | -0.0894 | -0.3398 | 0.3462 | -0.0885 | -0.1103 |
| BMI | -0.0894 | -0.0894 | 1 | -0.0088 | -0.0269 | 0.2524 | 0.0441 |
| ADHD | -0.1240 | -0.3398 | -0.0088 | 1 | -0.0902 | 0.0820 | 0.1188 |
| EDU PS | 0.2909 | 0.3462 | -0.0269 | -0.0902 | 1 | -0.1856 | -0.1865 |
| BMI PS | -0.0839 | -0.0885 | 0.2524 | 0.0820 | -0.1856 | 1 | 0.1413 |
| ADHD PS | -0.0630 | -0.1103 | 0.0441 | 0.1188 | -0.1865 | 0.1413 | 1 |

Note. Study variables were residualised for sex and 10 PCAs before computing the correlation table. GCSE scores were also residualised for age at obtention of GCSE. ADHD and BMI were a composite cross ages and were not residualised for age at data collection.

#### Section 2. Estimating the effect of X on Y in the single polygenic score case, based on observed associations

For the single polygenic score case depicted in Figure 3a, we assume the following structural equation model relating variables $G$, $X$ and $Y$:

$G = G$ (1)

$X = \beta_{GX} G + E_X$ (2)

$Y = \beta_{GY} G + \beta_{XY} X + E_Y$ (3)

where $E_X$ and $E_Y$ are error terms assumed to be uncorrelated with $G$ and $X$.
We assume that $G$, $X$ and $Y$ are each standardised and denote their pairwise correlations by $r_{GX}$, $r_{GY}$ and $r_{XY}$. Our aim is to estimate $\beta_{XY}$, the standardised effect of $X$ on $Y$ conditional on $G$, given observed or postulated values of the pairwise correlations.

Using standard results of linear algebra, the conditional effects $\beta_{GY}$ and $\beta_{XY}$ are given by

$$\begin{bmatrix} \beta_{GY} \\ \beta_{XY} \end{bmatrix} = \begin{bmatrix} 1 & r_{GX} \\ r_{GX} & 1 \end{bmatrix}^{-1} \begin{bmatrix} r_{GY} \\ r_{XY} \end{bmatrix}$$

Therefore the effect of interest is explicitly given by
$\beta_{XY} = \left(r_{XY} - r_{GX} r_{GY}\right) / \left(1 - r_{GX}^2\right)$ (4)

$\beta_{XY}$ can thus be derived from observed correlations, including those obtained from the best-fitting polygenic score.

In our sensitivity analyses depicted in Figure 5, we assume that polygenic score $G$ is a noisy measurement of a latent genetic value $G*$ that has the direct effects on $X$ and $Y$:

$$G = G^{*} + E_G$$

$$X = \beta_{G*X} G^{*} + E_X$$

$$Y = \beta_{G*Y} G^{*} + \beta_{XY} X + E_Y$$

We fit this latent variable model while constraining the variance in X explained by $G^{*}$ (in GsensX) or the variance in $Y$ explained by $G^{*}$ (in GsensY) to specified values, such as SNP- or twin-heritability. Specifically, $\beta_{G*X} = \sqrt{(h_X^2)}$ in GsensX, and $\beta_{G*Y} + \beta_{G*X}\beta_{XY} = \sqrt{(h_y^2)}$ in GsensY. In this case the observational correlations $r_{GX}$ and $r_{GY}$ decompose as $r_{GX} = r_{GG*} r_{G*X}$ and $r_{GY} = r_{GG*} r_{G*Y}$ respectively, and $r_{GX}/r_{GY} = r_{G*X}/r_{G*Y} = k$. The latent variable model thus estimates $k$ based on observed polygenic scores. However, the value of $k$ can be fixed if prior knowledge is available, such as $k = 0.5$ for mother-child pairs, as in Figure 1 and demonstrated in the Github repository.

#### Section 3. Complete genetic confounding in the single polygenic score case

Of note is that, based on $\beta_{XY}$ estimated above, it is possible to find what the heritability should be to completely confound an observed association, thereby rendering the conditional effect of X on Y null. From equation (4) we have:

$$\beta_{XY} = (r_{XY} - r_{GX} r_{GY}) / (1 - r_{GX}^2)$$

$$0 = (r_{XY} - r_{GX} r_{GY}) / (1 - r_{GX}^2)$$

$$r_{XY} = r_{GX} r_{GY}$$

Therefore, when the conditional effect of X on Y, $\beta_{XY}$ is null, then $r_{XY}$ is equal to the indirect path through G.

$r_{GY}$ is the square root of the heritability of Y, $r_{GY} = \sqrt{(h_y^2)}$

In the special case that X and Y are the same trait measured in a parent and child respectively, and assuming the heritability is constant across generations, then $r_{GX}$ is $0.5\sqrt{(h_y^2)}$ when G is measured in the child since its expected identity by descent with the parent is 0.5. Therefore $r_{XY} = 0.5 h_y^2$.

In this special case, the value of k is:

$$k = r_{GX} / r_{GY} = 0.5$$

#### Section 4. Estimating the effect of X on Y in the two polygenic scores case

For the two polygenic scores case depicted in Figure 4a the model equations are

$G_1 = G_1$ (5)

$G_2 = G_2$ (6)

$X = \beta_{G_1X} G_1 + \beta_{G_2X} G_2 + E_X$ (7)

$Y = \beta_{G_1Y} G_1 + \beta_{G_2Y} G_2 + \beta_{XY} X + E_Y$ (8)

and, under the same assumptions and notation as for the single polygenic score, the conditional effects are:

$$\begin{bmatrix} \beta_{G_1Y} \\ \beta_{G_2Y} \\ \beta_{XY} \end{bmatrix} = \begin{bmatrix} 1 & r_{G_1G_2} & r_{G_1X} \\ r_{G_1G_2} & 1 & r_{G_2X} \\ r_{G_1X} & r_{G_2X} & 1 \end{bmatrix}^{-1} \begin{bmatrix} r_{G_1Y} \\ r_{G_2Y} \\ r_{XY} \end{bmatrix}$$

in which $\beta_{XY}$ is the effect of interest.

In our sensitivity analyses depicted in Figure 7, we assume that polygenic scores $G_1$ and $G_2$ are noisy measurements of latent genetic values $G_1^*$ and $G_2^*$ that have the direct effects on $X$ and $Y$:

$G_1 = G_1^* + E_{G1}$

$G_2 = G_2^* + E_{G2}$

$X = \beta_{G_1^*X} G_1^* + \beta_{G_2^*X} G_2^* + E_X$

$Y = \beta_{G_1^*Y} G_1^* + \beta_{G_2^*Y} G_2^* + \beta_{XY} X + E_Y$

In GsensXY we fit this latent variable model while constraining the variances in $X$ and $Y$ explained by $G^*$ to specified values, such as SNP- or twin-heritability. Note that if these genetic variances equal the true heritabilities, the cross paths $\beta_{G_1^*Y}$ and $\beta_{G_2^*X}$ are zero; but otherwise they must be estimated in order to account for incomplete correlation between $G_1^*$ and $G_2^*$. However, we would generally expect the cross paths to have the same sign as the corresponding direct effects, i.e. $\beta_{G_1^*Y}$ and $\beta_{G_2^*Y}$ to have the same sign, and $\beta_{G_1^*X}$ and $\beta_{G_2^*X}$ also, assuming that $G_1^*$ and $G_2^*$ are positively correlated. When this is not the case, we refit the model with appropriate constraints on the signs of $\beta_{G_1^*Y}$ and $\beta_{G_2^*X}$.

### Annex A. Genetic overlap versus genetic confounding

This annex aims to clarify what we mean by 'genetic confounding' and how it differs from the concept of 'genetic overlap'. To help with this clarification, we rely on the model in Figure 3a and draw parallels between our notation and a common notation in the Mendelian randomization literature, i.e. $\beta$ is the direct effect of X on Y, $\gamma$ is the effect of G on X, and $\alpha$ is the direct effect of G on Y, corresponding to unmediated (also called horizontal) pleiotropy. As such, recalling that all variables are standardised, we have:

$r_{GX} = \gamma$ (9)

$r_{GY} = \alpha + \gamma\beta$ (10)

$r_{XY} = \beta + \gamma\alpha$ (11)

In (10) the observed correlation between G and Y is equal to the unmediated pleiotropy ($\alpha$) plus the pleiotropy mediated via the causal pathway ($\gamma\beta$). In (11) the observed correlation between X and Y is equal to the direct effect ($\beta$) plus the genetic confounding effect $\gamma\alpha$.

*Genetic overlap*

The genetic overlap comprises all of the association between X and Y linked to genetics, i.e. $r_{GX} r_{GY}$

Following the notation above:

$r_{GX} r_{GY} = \gamma\alpha + \gamma^2\beta$

The genetic overlap is thus decomposed into two components: the genetic confounding $\gamma\alpha$, and the direct effect $\beta$, scaled by the genetic variance of the exposure $\gamma^2$.

Subtracting the genetic overlap from the observed correlation yields:

$r_{XY} - r_{GX} r_{GY} = (\beta + \gamma\alpha) - (\gamma\alpha + \gamma^2\beta) = \beta - \gamma^2\beta$

Here, the residual association is the direct effect $\beta$ minus the component in $\beta$ that originates from G. The residual association can therefore be conceived as an ‘environmental association’ removing all effects coming from G, whether due to genetic confounding or to the effect of G mediated via the direct effect $\beta$.

*Genetic confounding*

Our expression for genetic confounding is:

$\beta_{GX} \beta_{GY}$

The difference from $r_{GX} r_{GY}$ is that $\beta_{GY}$ is conditional on X.

Similarly to equation (4) we can derive from the model equations (1-3) that $\beta_{GX} = r_{GX}$ and $\beta_{GY} = (r_{GY} - r_{XY} r_{GX}) / (1 - r_{GX}^2)$, so

$$\beta_{GX}\beta_{GY}=\hat{r}_{GX}\frac{\hat{r}_{GY}-\hat{r}_{XY}\hat{r}_{GX}}{1-\hat{r}_{GX}^{2}}$$

which, in the alternative notation can be written as:

$$\beta_{GX}\beta_{GY}=\gamma\left(\left(\alpha+\gamma\beta\right)-\left(\beta+\gamma\alpha\right)\gamma\right)/\left(1-\gamma^{2}\right)$$

Subtracting genetic confounding from the observed correlation yields:

$$r_{XY}-\beta_{GX}\beta_{GY}=\left(\beta+\gamma\alpha\right)-\gamma\left(\left(\alpha+\gamma\beta\right)-\left(\beta+\gamma\alpha\right)\gamma\right)/\left(1-\gamma^{2}\right)$$

which, after simplification is

$$r_{XY}-\beta_{GX}\beta_{GY}=\left(\beta+\gamma\alpha\right)-\gamma\alpha=\beta$$

The residual association here corresponds to the direct effect $\beta$.

In sum, we have two distinct sets of concepts:

1. Genetic overlap (capturing mediated + unmediated pleiotropy)
2. Genetic confounding (only unmediated pleiotropy)

And we have the two corresponding concepts for the residual associations:
1. Environmental Association, i.e. net of all genetic effects (which, in the absence of non-genetic confounders, would be lower than the causal effect)
2. Association net of genetic confounding (which, in the absence of non-genetic confounders would be equal to the  causal effect )
